# Why some size illusions affect grip aperture

**DOI:** 10.1101/866640

**Authors:** Jeroen B.J. Smeets, Erik Kleijn, Marlijn van der Meijden, Eli Brenner

## Abstract

There is an extensive literature debating whether perceived size is used to guide grasping. A possible reason for not using judged size is that using judged positions might lead to more precise movements. As this argument does not hold for small objects, and all studies showing an effect of the Ebbinghaus illusion on grasping used small objects, we hypothesized that size information is used for small objects but not for large ones. Using a modified diagonal illusion, we obtained an effect of about 10% on perceptual judgements, without an effect on grasping, irrespective of object size. We therefore reject our precision hypothesis. We discuss the results in the framework of grasping as moving digits to positions on an object. We conclude that the reported disagreement on the effect of illusions is because the Ebbinghaus illusion not only affects size, but –unlike most size illusions– also affects perceived positions.

---

Inspired by the two visual systems hypothesis (Goodale et al. 1991), an extensive literature has emerged on the question whether visual illusions affect our actions (Milner and Goodale 2006; Smeets and Brenner 2006; Franz and Gegenfurtner 2008). Sometimes illusions have an effect on action and sometimes they do not. Even within a single action, illusions frequently affect one aspect, leaving other aspects unaffected. For instance, using the Duncker illusion to change the apparent speed of a moving target that one is trying to intercept affects how quickly one moves to the target, but not where one aims to hit it (Smeets and Brenner 1995). Similarly, using the Ponzo illusion to change an object’s apparent size affects the way one lifts the object, but not the maximum grip aperture when reaching to grasp it (Brenner and Smeets 1996; Jackson and Shaw 2000). We have interpreted such findings as evidence that illusions only affect movement parameters that depend on the visual attribute that is affected by the illusion (Smeets et al. 2002). This interpretation does not explain *why* a movement parameter depends on a visual attribute. For grasping, our question therefore becomes: why do participants not use the size and position of the object but two positions on its surface to control the movements of the digits?

It has been proposed that we move in the way that we do in order to make our performance as precise as possible (Harris and Wolpert 1998), which means for many tasks that one moves in a way that minimizes the variance in the movement endpoints. Might the choice of using positions rather than size to control the digits be based on this leading to a better precision? When matching an object’s size with one’s digits, precision in reproducing the size with the distance between the digits decreases with increasing object size, whereas when moving to positions on the object’s surface, precision in the distance between the digits is independent of object size (Ganel et al. 2008a; Smeets and Brenner 2008). Smeets & Brenner (2008) provided a theoretical analysis of the measured precisions (Figure 1), and argued that “for objects that are larger than about 3 cm, relying on the positions of the object’s edges is more precise than relying on the object’s size”.

**Figure 1.**
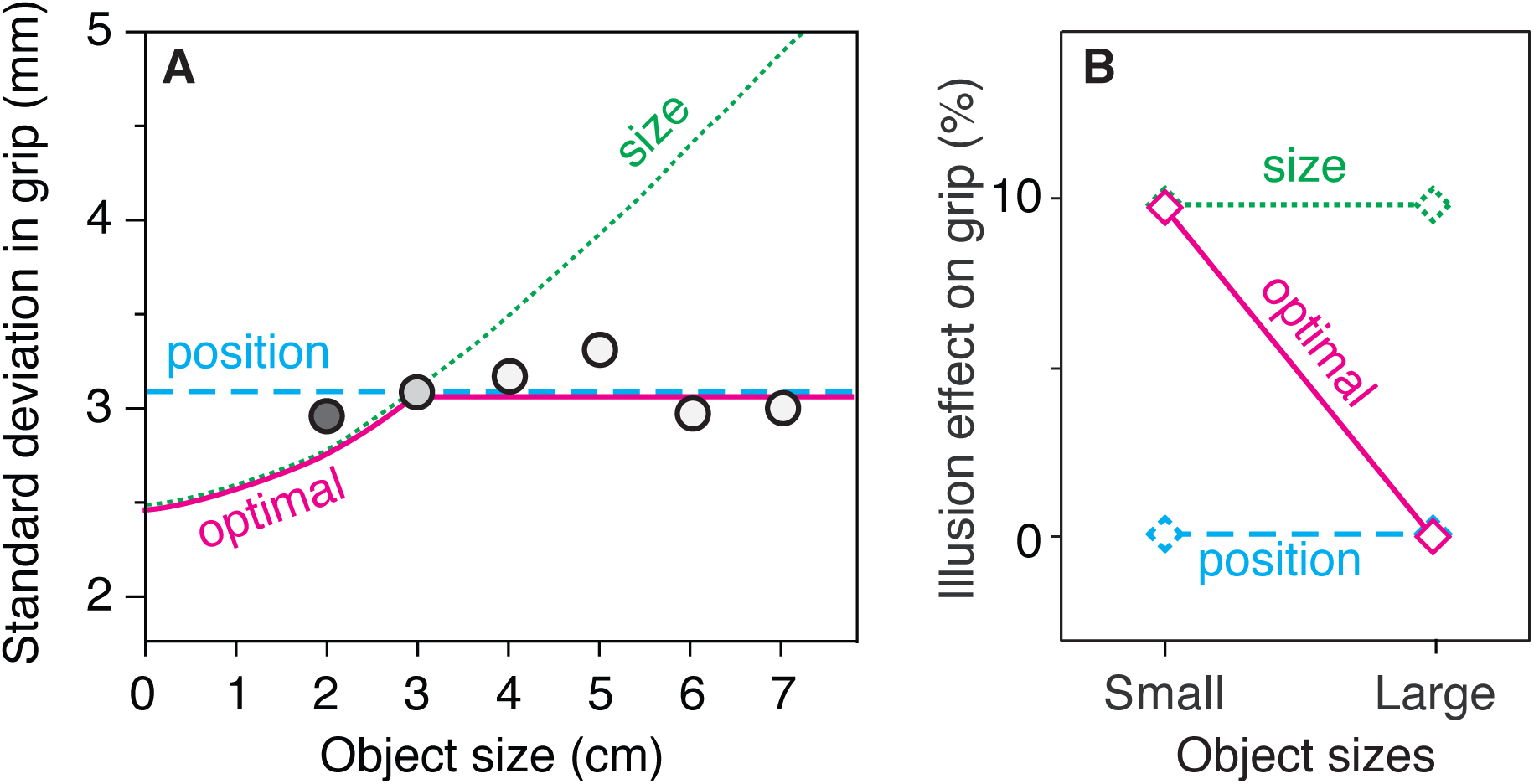
Model predictions. **A**) Possible interpretation of the data (dots) of Ganel et al. (2008a), based on the calculations by Smeets & Brenner (2008). The precision in maximum grip aperture would depend differently on object size if grip aperture depended on judgments of ‘size’ (green dotted curve) than if it depended on judgments of ‘position’ (cyan dashed curve). For each object size, the ‘optimal’ attribute to rely on (purple solid curve) is the lower (most precise) of the two. B) Prediction of the illusion effect on maximum grip aperture for small and large objects assuming a 10% effect of the illusion on perceived size.

Based on the analysis in Figure 1, one might argue that it would be optimal for participants to choose different information to guide the way they grasp objects of different sizes. They should guide their digits towards contact positions for large objects, but switch to scaling grip aperture to object size for objects that are smaller than 3 cm. Is there any evidence for this such an optimal precision hypothesis? Unfortunately, Ganel et al. (2008a) only had one object that was smaller than 3 cm (the dark-grey data point in Figure 1). The precision of grasping that object was between the ‘position’ and the ‘optimal’ predictions. In a similar experiment, Bruno et al. (2016) did have several objects that were smaller than 3 cm, and found that the variability in grip aperture increased with object size for these objects (but not for larger ones), which would be consistent with participants using judgments of size to guide their digits for the small objects. Such a trend is not visible in the data of a third study in which standard deviations in grip aperture were reported for various object sizes (Pettypiece et al. 2010), but this is not surprising given the lack of precision of the estimates in that study, as indicated by the large scatter around the fitted lines.

Showing that the choice of attribute that is used for controlling grasping depends on object size could solve a long-standing conflict about the question whether illusions affect maximum grip aperture. The two original studies that did not find an effect of the Ponzo illusion on maximum grip aperture both used relatively large objects (6.0-7.7 cm diameter; Brenner and Smeets 1996; Jackson and Shaw 2000). More recent studies in which no effect of illusions on maximum grip aperture were found also used objects that were larger than 3 cm: the Ponzo illusion (4-4.2 cm; Ganel et al. 2008b), the diagonal illusion (5.0-9.3 cm; Stottinger et al. 2009) and the empty space illusion (6-7 cm; Stottinger et al. 2012). On the other hand, in experiments that involved the Ebbinghaus illusion, authors generally used target objects with a diameter of about 3.0 cm (Aglioti et al. 1995; Haffenden and Goodale 1998; Pavani et al. 1999; Franz et al. 2000; Haffenden et al. 2001; Kopiske et al. 2016). In these experiments, a consistent effect of the illusion on grip aperture is reported (Franz and Gegenfurtner 2008), although whether the effect is as large as that on perception is still under debate (Kopiske et al. 2017; Whitwell and Goodale 2017). So, the optimal precision hypothesis can explain many experimental findings: size illusions only affect maximum grip aperture if object diameter is 3 cm or smaller.

There is an alternative explanation for the experiments using the Ebbinghaus illusion showing a clear effect of perceived size on maximum grip aperture, whereas most experiments using other illusions don’t. The basis of this alternative explanation is the digit-in-space hypothesis: grasping is always based on moving one’s digits to locations in space, rather than shaping one’s hand opening to object size (Smeets and Brenner 1999; Smeets et al. 2019). According to this hypothesis, maximum grip aperture has nothing to do with judged size, but is based on judged positions. Given the finding that the Ebbinghaus illusion, unlike other illusions, influences the judged positions of points on the object’s surface (Smeets and Brenner 2019), one expects maximum grip aperture to be affected by the Ebbinghaus illusion, but not by other size illusions. This explanation is irrespective of whether objects are small or large.

We therefore decided to investigate whether an illusion that does not affect perceived positions influences maximum grip aperture for objects of different sizes. We used the modified diagonal illusion, because its effect on the perceived size of a single object is similar to that of the Ebbinghaus illusion, without any effect on the perceived positions of locations on the object’s edges (Smeets and Brenner 2019). Moreover, this illusion does not rely on surrounding objects that might influence grip aperture because they are regarded as obstacles (Haffenden et al. 2001; de Grave et al. 2005; Biegstraaten et al. 2007). The prediction of the optimal precision hypothesis is that this illusion will not affect maximum grip aperture for large objects (> 3 cm), but for small objects (< 3 cm) the effect will be comparable to the perceptual illusion (indicated by the solid purple line in Figure 1B). The alternative explanation based on the digit-in-space hypothesis predicts that there will be no illusion effect on maximum grip aperture, irrespective of object size (cyan dashed line in Figure 1B). A third possibility, based on the premise that grip aperture is always based on the perceived size (Franz 2001; Kopiske et al. 2016), is that the illusion will influence maximum grip aperture as much as it does perception, irrespective of object size (green dotted line in Figure 1B).

## Methods

### Participants

Nineteen right-handed volunteers (age range 18-60, mean 30 years) participated in the experiment. Assuming that the standard deviation in the effect of an illusion on maximum grip aperture is about 1.8 mm (Kopiske et al. 2016) and the illusion effect is about 1.5 mm (5%; Smeets and Brenner 2019) we should obtain a power of about 0.98 with 19 participants. All participants had normal or corrected to normal vision and were naïve with respect to the hypothesis that was tested.

### Stimuli and equipment

In choosing the illusion, we took care to ensure that the illusion-inducing context did not introduce structures that could be regarded as obstacles, that the illusion is not based on a contrast between two simultaneously presented targets, and that the effect of the illusion-inducing context was robust for small objects. To achieve this, we combined elements of two illusions: the diagonal illusion and the empty space illusion. The empty space illusion is the phenomenon that a space filled with many elements seems larger than a space that is empty or contains a single element (Luckiesh 1922; Stottinger et al. 2012). We combined this illusion with the diagonal or Sander illusion (Luckiesh 1922; Stottinger and Perner 2006) to produce a modified diagonal illusion. In pilot experiments, we determined parameters of the stimulus that were effective in inducing illusory length differences. The same parameters have been used in an experiment on the perceptual effects of this illusion (Smeets and Brenner 2019).

As consistent haptic feedback is essential for normal grasping (Cuijpers et al. 2008; Schenk 2012), we let our participants grasp real wooden bars that had a 4×4 mm section. We had two short bars (1.5 & 2.5 cm) and two long bars (4 & 5 cm). We used two bar lengths within each category in order to ensure that participants process the visual information of the stimuli and do not simply categorize the stimuli as either ‘short’ or ‘long’ and use learned responses. For each length, we had one bar with a single white dot and another bar with several (3-5) dots. We presented each bar on a sheet of paper with a quadrilateral printed on it. There was a different sheet of paper on the table for each bar, so that the influences of the number of dots and of the kind of quadrilateral enhanced each other (Figure 2). The bar (and thus quadrilateral) was always at the same position in front of the participant, oriented sagittally.

We instructed the participants to start their movements grasping a 40 mm long, horizontal rod, positioned 30 cm to the right of the target. When grasping this rod they had a grip aperture of 4 mm. When starting in this way, the hand’s movement is perpendicular to the bar, which not only makes grasping easy, but also ensures that the hand does not cover the illusion (Carey 2001). We deliberately chose to make the target continuously visible, because the predicted availability of feedback at contact influences how grasping movements are planned (Bozzacchi et al. 2018). Continuous vision of the target furthermore ensures direct visuomotor control without memory effects (Gentilucci et al. 1996; Westwood and Goodale 2003) and a proper visual-proprioceptive integration of location information (Smeets et al. 2006). In order to prevent participants from adjusting grip aperture on the basis of a direct visual comparison between target and digits (Haffenden and Goodale 1998; Franz et al. 2001; Bruno and Franz 2009), we placed a horizontal wooden board above the table, covering 80% of the trajectory of the hand towards the target.

**Figure 2:**
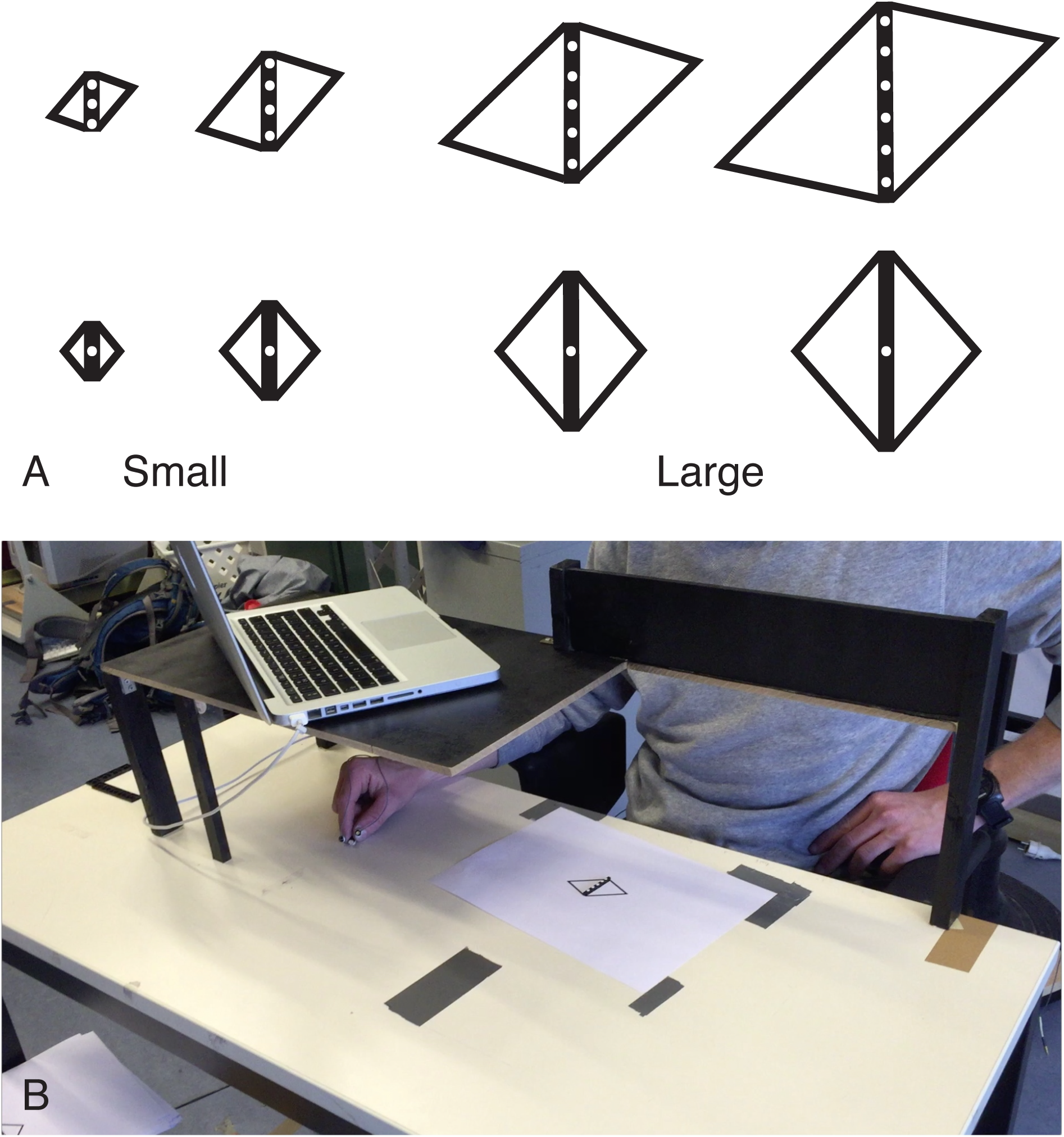
Methods. **A**) The eight stimuli used in the experiment. **B**) A participant in the set-up with his hand holding the starting bar.

The illusion works best when viewed from above, so we placed a 10 x 40 cm board vertically 20 cm above the table in front of the participant, 13 cm closer to the participant than the bar. This board forced participants to move their head forward in order to be able to see the target. In this way, we ensured that participants viewed the bar from above both when matching and grasping, and that they could not see the bar between trials. We used an Optotrak 3020 (Northern Digital) to record the movements of markers attached to the nails of finger and thumb at 200 Hz. For the perceptual judgements, we placed a laptop on the wooden board, above the start location of the hand. This laptop showed a black screen (28.5×18 cm; surrounded by a 1 cm thick black frame) with a 2mm thick vertical white line presented at the centre of the screen; the length of the line could be adjusted by sliding one’s finger on the trackpad.

### Tasks and procedures

In order to ensure that participants looked at the bar before starting to move their hand, participants always reported the perceived size of the object directly before they grasped it. This previewing may have increased the influence that perceptual judgements of size have on the grasping movement (Glover 2002), so the influence on grasping might be overestimated.

Each trial started with the experimenter placing a stimulus (combination of a bar and a piece of paper with a drawing of the corresponding context, Figure 2). A line with random length was then presented on the screen of the laptop and participants moved their hand to the trackpad to match the length of the line on the screen to that of the bar. After clicking the trackpad to confirm the match, participants moved their right hand to the starting bar. They were then given a go-signal, indicating that they should pick up the bar in a single continuous movement, and place it at the target area (indicated by the thick black piece of tape on the table). This resulted in reach-to-grasp movements with a duration of about 600 ms. After placing the bar on the table participants moved their hand back to the starting position, and the next trial began.

Every stimulus was presented 10 times in a pseudorandom sequence, so in total there were 80 trials, each consisting of a judgement and a grasping movement. If the participant dropped the bar before reaching the end position, or if the bar was not grasped at the ends with a precision grip, we repeated the grasping part of the trial. After every 20 trials, the participants were asked whether they wanted a short break.

### Data analysis

We analysed the kinematic data of each grasping movement. We determined the grip aperture by taking the 3D distance between the markers of finger and thumb. We subsequently determined the maximum of this distance (MGA) in the part of the trajectory between the start and contact. Contact was determined as the first moment after peak height that both digits were less than 1 cm above the level of their starting position. We checked by visual inspection of all trials that this method yielded a sensible measure (e.g. that MGA was not affected by repositioning the digits after initial contact).

To determine the illusion effect, we started by taking the median response (matched length or MGA) for each participant, stimulus and task. The advantage of using the median rather than the mean is that it is robust for outliers. The difference between such median values for the two illusion configurations for the two stimuli containing an object of the same size provides us with 4 raw illusion effects for each participant. These effects were scaled by dividing them by the slope of a fit between the eight median values and the corresponding actual object sizes (Glover and Dixon 2001; Franz et al. 2005; Hesse et al. 2016). We subsequently expressed the illusion effects as a percentage of the mean matched length or MGA, and averaged these percentages for the two small and for the two large objects. All this was done separately for each participant.

We analysed the resulting average percentage data with a repeated measures ANOVA with factors size (small, large), and task (matching, grasping). The precision hypothesis primarily predicts an interaction, because it predicts that the illusion effect on matched length will be the same for both sizes (green dotted symbols in Figure 1b), whereas the illusion effect on MGA will close to zero for large objects (solid purple line). The alternative explanation based on the digit-in-space hypothesis predicts only a main effect of task, because it predicts the same illusion effect for the matching task as the precision hypothesis, combined with no illusion effect on grasping for both sizes (cyan dashed line in Figure 1b).

In an additional analysis, we examined whether our data produce the pattern of variability within each condition that was the basis of the precision hypothesis. We tested whether objects that were perceived as being larger were matched with more variability. We also examined to what extent the variability in grip aperture depends on the grip aperture. For this, we determined the standard deviation in the maximum grip aperture and in the matching response for each participant and each of the four object sizes and two illusion configurations. We analysed both measures (variability in matched lengths and in MGA) in separate 4 (size) x 2 (illusion) repeated measure ANOVAs.

As a second additional analysis, we determined whether the illusion effects are correlated across participants. If the precision hypothesis is correct, we expect the illusion effects for MGA to be correlated with the effects for matching for the small objects, but not for the large objects. If grip aperture is always based on the perceived size, we expect the illusion effects for MGA to be correlated with the effects for matching for both object sizes. A similar correlation between the effect of the illusion on MGA for small and large objects is expected according to all hypotheses except the optimal precision hypothesis, according to which there should be no correlation. No correlation is expected if grip aperture is based on judged positions. Correlations are quite difficult to determine for this type of data (Franz et al. 2001). To make sure that the absence of a correlation is not due to a lack of power, we not only determined the correlations that are predicted by the various hypotheses, but also a correlation that should be present irrespective of the hypothesis that is correct: that between the perceptual effect of the illusion for small and large objects.

### Results

We designed our experiment to test predictions about the effects of a size illusion on maximum grip aperture. We have three mutually excluding predictions for these effects, expressed relative to the effect of the illusion effect on the matching task (Figure 1B). The first one is that if <u>*size*</u> is used to control grip aperture, the mean effect of the illusion on maximum grip aperture will be the same as for matching, irrespective of object size. The second one is that if <u>*positions*</u> are used to control grip aperture, the illusion will have no effect on maximum grip aperture, irrespective of object size. The third is that if the information with the highest precision is used to achieve an <u>*optimal*</u> performance, as explained in the introduction, the effect of the illusion on maximum grip aperture will be the same as for matching for small objects, but there will be no effect for large objects.

The results are very clear (Figure 3A): for both the large and the small objects, there is a considerable illusion effect for perceptual matching, and no illusion effect on maximum grip aperture. This pattern is confirmed by the ANOVA: a main effect of task (F_1,18_ = 50.2; p<0.01) with neither an effect of size (F_1,18_ = 1.05; p=0.32) nor an interaction (F_1,18_ = 0.53; p=0.48). As there is no interaction between task and size (the non-significant slightly larger effect of the illusion when grasping larger objects is even in the opposite direction than predicted), we can reject our precision hypothesis (solid purple line in Figure 1B). As the illusion effect on maximum grip aperture does not differ from zero, either for small or for large objects (see confidence intervals of the open symbols), the results are in conflict with the predictions based on the use of size, and in line with the predictions based on using positions for controlling grip aperture.

**Figure 3.**
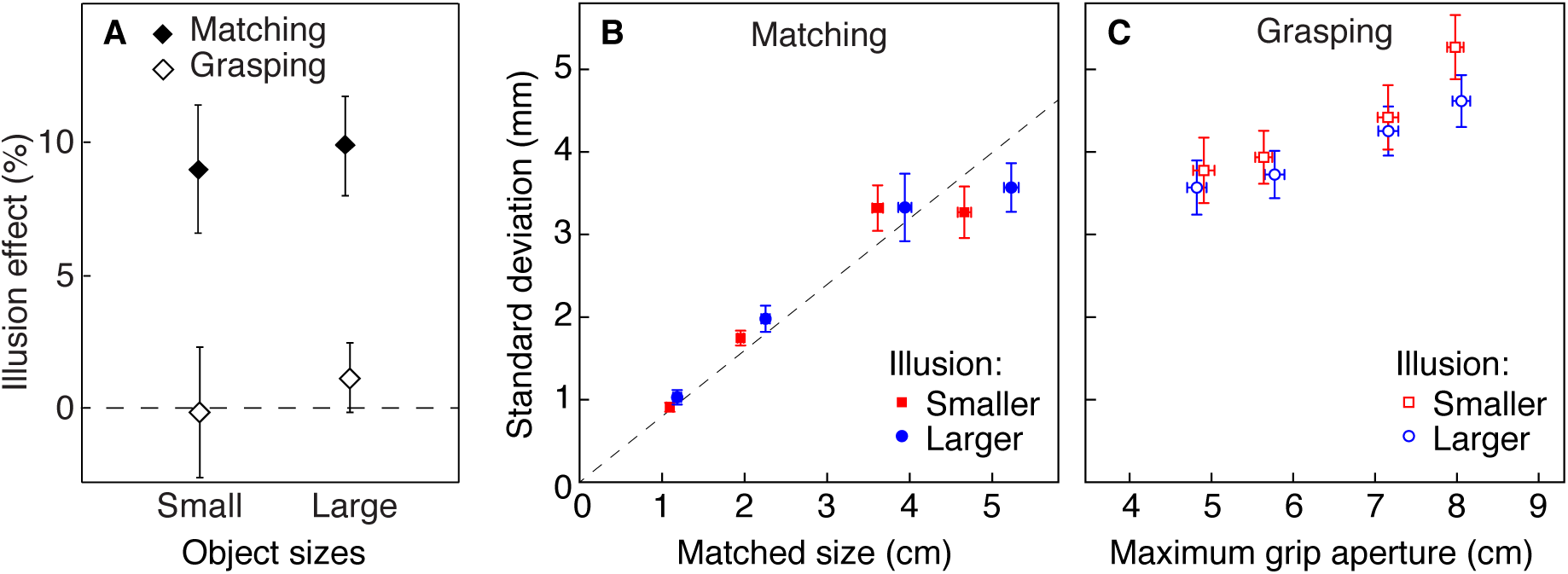
Results. All values are averages with 95% confidence intervals across participants. **A**) The illusion effects for the two tasks and object size categories (small:1.5 and 2.5 cm; large: 4.0 and 5.0 cm). **B**) Within participant standard deviation in matched size as a function of matched size for each of the four objects in the two illusion-inducing contexts. The dashed black line shows a Weber fraction of 8%. **C**) Within participant standard deviation in maximum grip aperture as a function of maximum grip aperture.

We checked whether the pattern of variability corresponds to the pattern reported by Ganel et al. (2008a). For the perceptual match (Figure 3B), we found a strong increase in variability with object size (main effect in the ANOVA: F_3,54_ = 47.0, p<0.001). The variability in the matched size was three times as high for the largest object than for the smallest one. There was a small effect of the illusion (F_1,18_ = 4.67; p=0.044), with no interaction with size (p=0.81). Figure 3B shows that the small effect of the illusion on the variability follows the illusion’s effect on the mean matched size: the larger appearing bars were matched with more variability than the smaller appearing ones (blue dots are to the right and above the corresponding red squares). The results can be described well with a Weber fraction of 0.08 in matching performance (dashed line), which is slightly less precise than the value of 0.06 that we used for the predictions in Figure 1 (Ganel et al. 2008a; Smeets and Brenner 2008).

The maximum grip aperture increased in a normal way with object size: it ranged from 4.8 cm for the 1.5 cm object to 8 cm for the 5 cm object. Its standard deviation was on average 4.2 mm (Figure 3C), so grip aperture was more variable than it was in the data of Ganel et al (2008a), close to the value that was reported by Bruno et al. (2016). We found a moderate increase of variability with object size (slope 0.03; F_3,54_ = 12.5, p<0.001), without a significant effect of illusion (p=0.13) or interaction (p=0.68). We will come back to this finding in the discussion. The non-significant tendency for smaller appearing objects to be grasped with a more variable maximum grip aperture than larger appearing ones (red squares higher than blue circles in Figure 3C) is in the opposite direction than the significant effect of the illusion that we found in the variability in perceptual matching (blue dots higher than red squares in Figure 3B).

We find no correlation between the illusion effects for matching and maximum grip aperture for either of the bar sizes (both p>0.7; upper row in Figure 4). This is inconsistent with any hypothesis that involves judged size being used to control grip aperture. It is consistent with the hypothesis that positions are used to control grip aperture. We find significant positive correlations between the illusion effects on maximum grip aperture for small and large objects as well as between the illusion effects on matching for small and large objects (both p<0.05; lower row in Figure 4). The latter findings indicate that the illusion effects are determined reliably enough to give our experiment the power that is needed to detect correlations within the data with this number of participants. Furthermore, the correlation that was found for grip aperture adds to the evidence against our optimal precision hypothesis.

**Figure 4.**
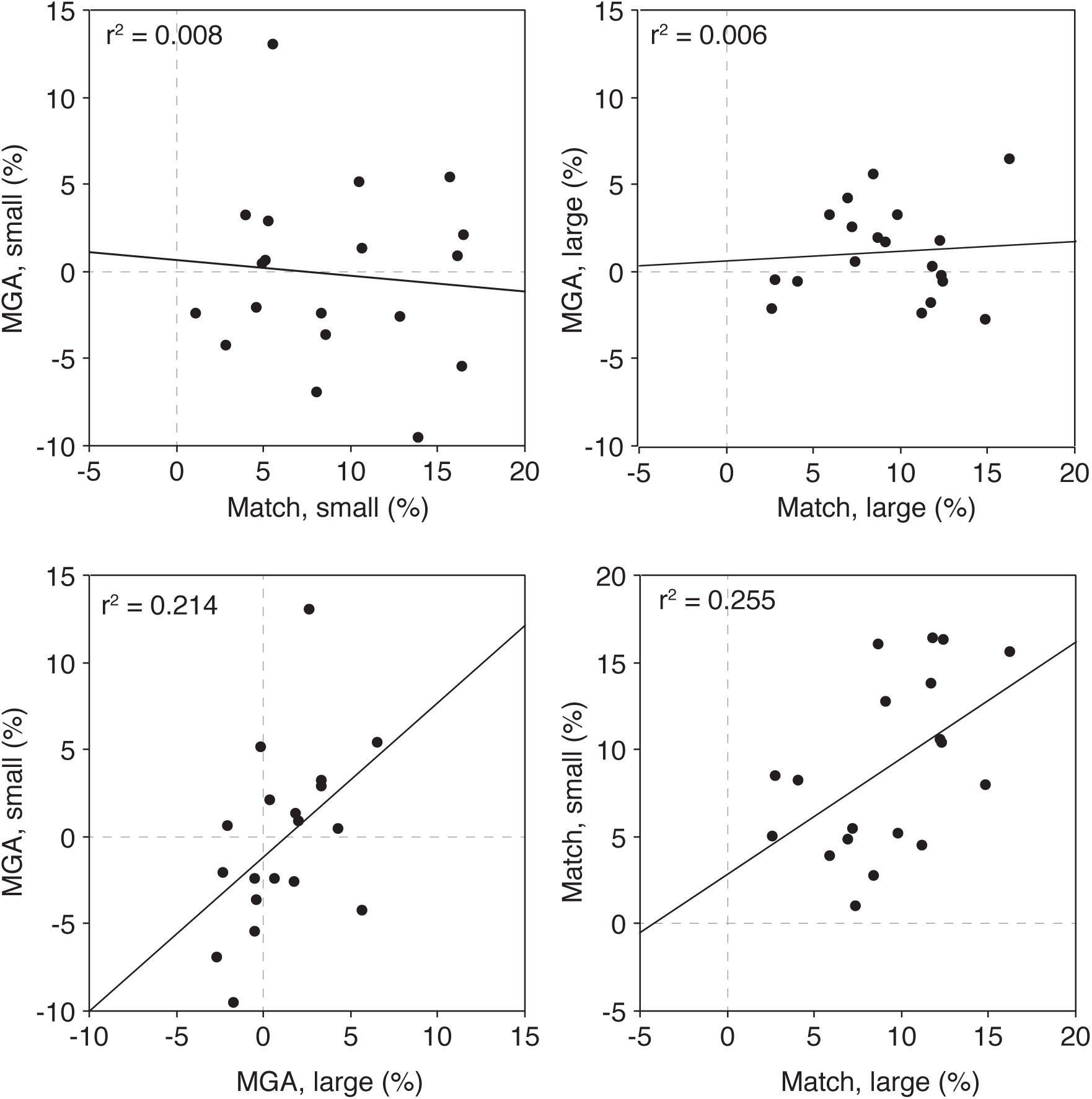
Correlations between the illusion effects across tasks and sizes. Each symbol represents the average value for a single participant. The numbers that are reported (r^2^) are squared Pearson’s correlation coefficients. Upper row: the illusion effect on maximum grip aperture is not correlated with the illusion effect on matching. Lower row: the illusion effect on small objects is positively correlated with the illusion effect on large objects within both tasks.

## Discussion

The data very convincingly reject the optimal precision hypothesis: the maximum grip aperture was not affected by the illusion for any object size (open diamonds in Figure 3A correspond to dashed cyan line in Figure 1B). Our data also reject the hypothesis that grip aperture is based on perceived size (green dotted line in Figure 1B): in our experiment the illusion effect on maximum grip aperture is clearly different from that on perceptual matching (solid and open symbols differ in Figure 3A). These two conclusions are supported by the correlation analysis: the variations between participants in the (on average absent) effect on grip aperture are not correlated with the variations in effect on perceptual matching, irrespective of object size, whereas the variations are correlated between the small and large objects (Figure 4). The results are consistent with the hypothesis that participants always use position information to guide their digits during grip formation. This suggests that they use a globally optimal strategy in grasping, rather than a strategy that is optimal for the specific size of the object that is to be grasped.

Assuming that the aim of grasping movements is to bring digits to their contact points on the object (Smeets et al. 2019), it is evident that information about such contact points is essential for a normal control of grasping movements. It is therefore not surprising that withholding information about the contact points or how they can be reached can influence the whole grasping movement. This has been demonstrated for haptic information (Cuijpers et al. 2008; Schenk 2012; Davarpanah Jazi and Heath 2016) as well as for visual information (Whitwell et al. 2016; Bozzacchi et al. 2018). We therefore designed our set-up in a manner that provided our participants with both visual and haptic feedback near the point of contact, despite restricting the visual feedback to the last part of the digits’ trajectories. In this way, participants could use feedback to keep their movements calibrated, without being able to use feedback to correct them on-line.

The availability of visual information near contact is important in the study of the effect of illusions on motor control. It has been shown that the Brentano illusion has a larger effect on pointing if visual feedback is removed, which can be interpreted as a shift from relying on positions to relying on sizes (de Grave et al. 2004). The reduced effect of the illusion when vision was available was not due to on-line movement corrections when close to the target, because it was independent of movement speed. A possible explanation is that egocentric position information deteriorates more quickly, so it is advantageous to shift to using size when vision is removed. In grasping, a similar shift away from using positions might happen if vision is blocked. This might explain why an open-loop grasping experiment in which all vision was blocked once the hand started to move found a clear effect of the diagonal illusion on maximum grip aperture (Whitwell et al. 2018).

In our experiment, the peak in grip aperture occurred when the hand was already in view. This raises the possibility that seeing the hand near the target could have led to on-line corrections that cancelled an illusion effect. We think this is unlikely because the hand was visible for less than 200 ms before reaching maximum grip aperture for all participants, whereas it takes more than 200 ms to complete any on-line correction (Oostwoud Wijdenes et al. 2011). Nevertheless, we checked whether there were clear effects of the illusion earlier in the movement by determining the effect of the illusion on grip aperture at 2/3 of the movement: when the digits were 20 cm from the starting position. At that position, the digits are still invisible. If the illusion would have influenced grip aperture in accordance with its effect on perceived size, we would expect an illusion effect of about 2 mm at that position (2/3 of the 3 mm perceptual effect). We did not find any illusion effect on the grip aperture: the mean effect is 0.3 mm, with a standard error across participants of 1.3 mm. Thus, the lack of illusion effect on maximal grip aperture is not due to on-line corrections based on visual feedback during the last part of the trajectory.

Our hypothesis was based on published data on the precision of grip aperture during grasping and size matching (Ganel et al. 2008a). Our data show roughly the same pattern, but with a slight difference: we reproduce the clear increase in variability with object size for the matching task, but we also see a modest, significant increase in variability for maximum grip aperture (Figure 3C). When re-examining the data of Ganel et al. (2008a), we see a similar increase in variability (the leftmost four data points in Figure 1A) with an even shallower slope than in our data. This increase is not always found: two recent experiments even report a slight decrease of variability with object size for the range of sizes that we tested (Utz et al. 2015; Bruno et al. 2016). Altogether, there is therefore no consistent change in the variability in maximum grip aperture with object size. Whether the variability is completely independent of object size is not relevant for our conclusion, because in all experiments, the variability increased much less than was to be expected on the basis of Weber’s law for size.

The variability within participants varies considerably between studies. Our participants were about as variable in reproducing maximum grip aperture as were those of Bruno et al. (2016) and Pettypiece et al. (2010), but were much less variable than those of Utz et al. (2015). Our participants were about 30% more variable than the participants in the study of Ganel et al. (2008a), both in terms of the maximum grip aperture and in their size judgements. There are various differences between the experiments that might explain the larger variability. The most likely explanation of the 30% larger variability in our study is the larger viewing distance (more than 40 cm in our experiment versus 30 cm for Ganel et al. 2008a). A second likely origin of the larger variability in our experiment is that we alternated between the two tasks, whereas Ganel et al used a blocked design. A third factor that might have played a role is the difference between the viewing conditions (Desmurget et al. 1997): Ganel’s participants could see their hand together with the target before movement onset, whereas our participants did not see their hand until it was close to the target object. As grip aperture not only depends on the present trial, but also on what happened on the previous trial (Tang et al. 2015), a fourth factor that is important for variability is the range of sizes. However, as we used a smaller range than Ganel et al. (2008a), this factor would predict less variability in our study. The shape of a target is also important for the way one opens one’s hand in grasping (Verheij et al. 2012; Verheij et al. 2014), but as both studies used thin bars, this cannot be the basis of the difference.

The strength of the illusion that we found here (about 10%) is larger than in our pencil-and-paper version of the experiment (5%; Smeets and Brenner 2019). There are several differences between the studies that might account for this difference. The main difference is the viewing geometry. In the present experiment, the bar was oriented in the sagittal direction and viewed from above. In our pencil-and-paper experiment, the bar was oriented in the fronto-parallel direction and the viewing angle was unconstrained.

A recent extensive experimental paper demonstrated convincingly that grasping is affected by the Ebbinghaus illusion, and that this effect was not due to the flankers acting as obstacles (Kopiske et al. 2016). How can we reconcile their clear effect of the illusion on grip aperture (comparable in size to the perceptual effect) with the total absence of an illusion effect on maximum grip aperture in the current study? Our pencil-and-paper experiment showed that this discrepancy is actually consistent with assuming that grip aperture is based on position perception (Smeets and Brenner 1999; Smeets et al. 2019): the Ebbinghaus illusion does not only affect perceived size, but also perceived positions, whereas the modified diagonal illusion only affects perceived size. We have no idea why the illusions have such different effects on perceived positions while influencing perceived size in a similar manner, but a somewhat similar distinction has been observed for orientation illusions (Dyde and Milner 2002).

The present study provides a comprehensive explanation for the apparently conflicting results on the effects of visual size illusions on grip aperture in reach-to-grasp movements. Size illusions themselves have no effect. The position illusion that is present in the Ebbinghaus figure is responsible for the robust effect of that ‘size’ illusion.

## Supplementary Material

The data for grip aperture and perceptual settings for each trial and subject are available at http://osf.io/y22jj

